# Analysis of D- and L- Isomers of (Meth)amphetamine in Human K2EDTA Plasma

**DOI:** 10.1101/2022.11.20.517241

**Authors:** Brian Robbins, Jacob Perry, Mary Long, Rob E. Carpenter

## Abstract

Methamphetamine and its metabolite amphetamine are frequently abused drugs. Whether obtained legally or from clandestine laboratories it is of relevance to determine the chiral makeup of these drugs for investigative purpose. Although urine and oral fluid matrices are commonly offered, less available to independent laboratories are techniques to verify dextro (D-) or levo (L-) (meth)amphetamine from human K2EDTA plasma. This paper outlines the development and validation of a method that includes the addition of internal standard and a two-step liquid-liquid extraction to remove the analytes from human K2EDTA plasma by triple quadrupole mass spectrometry (LC-MS/MS). The assay was validated according to the United States Food and Drug Administration and College of American Pathologists guidelines, including assessment of the following parameters in plasma validation samples: linear range, limit of detection, lower limit of quantitation, matrix effects, inter- and intra-day assay precision and accuracy, carry over, linearity of dilution, matrix effects and stability. The outcome is a validated and reliable method for the determination of D- and L-isomer concentration of meth(amphetamine) human plasma samples that can be easily adopted by independent clinical laboratories.

## 1. Introduction

With the rising recognition that drugs of abuse increased during the COVID-19 pandemic [1], independent clinical laboratories are called upon, now more than ever, to offer testing solutions for diagnostic and investigative purpose. Methamphetamine is a central nervous system (CNS) stimulant whose prevalence has shown significant increase in (mis)use during recent years [2–4]. Importantly, methamphetamine and its metabolite amphetamine exist as two enantiomeric forms, dextro (D-) or levo (L-), which produce radically different effects on the CNS [5]. Aside from the potential use disorder from the dopamine response that (meth)amphetamine asserts [5], therapeutic doses of each enantiomer are commonly prescribed for ADHD, narcolepsy, and severe obesity [6,7]. Furthermore, the L-enantiomer is an effective vasoconstrictor used in the over-the-counter formulation of Vicks Vapor Inhaler [6]. This can be problematic when determining the source of (meth)amphetamine in patient samples that can be easily adulterated.

Although liquid chromatography tandem mass spectrometry (LC-MS/MS) assays that detect (meth)amphetamine from human urine or oral fluid matrices are commonplace, specificity of D- and L- isomers can be methodological challenging [8,9]. And although human blood plasms LC-MS/MS assays that identify (meth)amphetamine have been deployed [10], less available are plasma assays for accurate enantiomeric delineation for independent clinical laboratories. Accordingly, this study describes a validated laboratory develop LC-MS/MS assay to quantify the enantiomeric forms of (meth)amphetamine) in human blood plasma samples. The assay was developed in parallel with an assay for 63 electro spray (ES) positive analytes that reflected identify D-- and Lforms of (meth)amphetamine [11]. The result is a human blood plasma assay that provides an accurate quantitative view of on-board D- and L- enantiomeric (meth)amphetamine in the blood stream for the independent clinical laboratory.

## 2. Materials and Methods

### 2.1. Reagents and Standards

All analyte stock solutions at 1 mg/mL concentrations and deuterated internal standards at 100 µg/mL were purchased from Cerilliant Corporation (Round Rock, TX, USA). All organic solvents including methanol, acetonitrile, formic acid (88%), dichloromethane, 2 propanol and ethyl acetate were obtained from Fisher Scientific (Pittsburgh, PA, USA). Blood plasm was collected with the VACUETTE® K2 DTA Blood Collection Tube.

### 2.2. Mobile Phase and Extraction Solutions

A D- and Lmobile phase (MPDL) solution was created by adding ∼993.2 mL of methanol to a 1L bottle. Then using a pipettor, 5 mL of type I clinical grade water, 1.5 mL of acetic acid, and 0.3 mL of ammonium hydroxide were added. This solution can be kept at room temperature for up to 1 year. Mobile phase A was prepared by adding 974 mL of LCMS grade deionized water, 25 mL of methanol, and 1 mL of 88% formic acid to a clean 1 L reagent bottle which was capped and mixed well. This solution can be stored at room temperature for up to 2 weeks. Mobile phase B was prepared by adding equal parts methanol and acetonitrile to a clean reagent bottle which was capped and mixed well. This solution can be kept at room temperature for up to 1 year. Extraction solution 1 (ES1) was created with 50% dichloromethane and 50% 2-propanol by using a graduated cylinder under a fume hood. Equal volumes of dichloromethane and 2-propanol were added to a clean reagent bottle which was capped and mixed well. Extraction solution 2 (ES2) was created with 50% dichloromethane and 50% ethyl acetate by using a graduated cylinder under a fume hood. Equal volumes of dichloromethane and ethyl acetate were added to a clean reagent bottle which was capped and mixed well. ES1 and ES2 can be kept at room temperature for up to 1 year.

### 2.3. Standard Preparation

An 8000 ng/mL stock solution was made by combining analyte stock controls and diluting it with mobile phase A (MPA). In contrast, D- and Lamphetamine and methamphetamine were added in an amount to make a 4000 ng/mL stock of each isomer so that combined they would produce an 8000 ng/mL solution of total amphetamine and methamphetamine. This means that the range of the D- and Lstandard curve (SC) is from 2.5 to 1000 ng/mL (half the concentration). The resulting stock standard was diluted with MPA to produce the SC. Concentrations were 8000 (undiluted), 4000, 2000, 1000 400, 200, 100, 40, 20, 10, 4 and 2 ng/mL. The assay quality controls (QCs) were made similarly; first making a 7200 ng/mL spiking solution in MPA then diluting to 3200, 2400, 300, 60, 12, and 2 ng/mL. The D- and Lamphetamine and methamphetamine QCs were made at half concentrations and these solutions were stored at the concentrations above. During preparation the standards and quality controls were combined 1 part standard and 3 parts plasma to make a 250 µl sample volume. The final D- and LSC and QC concentrations were QC: 1000, 500, 250, 125, 50, 25, 12.5, 5, 2.5, 0.5 and 0.25 ng/ml and QC: 900, 400, 300, 37.5, 7.5, 1.5, and 0.25 ng/ml.

The internal standard working solution (ISWS) for the P63 assay and D- and L- assay was made by filling a 100 mL graduated cylinder to the 50 mL mark with 10% methanol in water and adding 250 µL of each of the internal standards listed above. The volume was brought to 100 mL with additional 10% methanol producing a concentration of 250 ng/mL.

### 2.4. Instrumentation

The liquid chromatography components of the LC-MS/MS system consisted of a model CBM-20A controller, 2 model Prominence LC-20AD pumps, a model DGU-20A5 degasser and a model SIL-20AC autosampler all obtained from (Shimadzu, Columbia MD, USA, based in Kyoto, Japan). The mass spectrometer used was a SCIEX API 4000 and the acquisition software was Analyst, v 1.5.2, build 5704 (Framingham, MA, USA). Nitrogen was obtained using a Peak ABN2ZA gas generator (Peak Scientific, Billerica, MA, USA). Reagents were weighed on a Mettler Toledo MX5 analytical micro balance (Fisher Scientific, Pittsburgh, PA, USA). Samples were dried on a TurboVap® LV (Uppsala, Sweden). Samples were vortexed on a Fisherbrand 120 multitube vortex. The analytical column was an Astec CHIROBIOTIC® V2 5.0 µm (2.1mm x 25 cm column) Catalog # 15020AST SUPLECO®, (Bellefonte, PA, USA).

### 2.5. Analyte Optimization

Individual analytes and internal standards were optimized by using T-infusion with 50%B mobile phase and tuning for declustering potential (DP), entrance potential (EP), collision energy (CE) and exit potential (CXP) at a flow rate of 0.7 mL/min. The two most abundant fragments were selected for monitoring using multiple reaction monitoring (MRM).

### 2.6. Sample Preparation and Procedures

The samples, standards and QC were extracted using two liquid-liquid extractions with 1:1 dichloromethane (DCM): isopropyl alcohol (IPA) and 1:1 DCM: ethyl acetate (EtAc). They were combined, dried, reconstituted with 1:1 methanol (MeOH): water and combined with mobile phase A for separation of the initial 63 analytes. Sample preparation for D- and L-analysis by LC-MS/MS involved transferring 50 µL of the already extracted standards, QC, and any samples of interest to a new plate. Then 450 µL of MPDL was added to each well and mixed with a multichannel pipette, the plate was covered with a plate mat and analyzed for the D- and L- isomers of amphetamine and methamphetamine using the listed chiral column. The LC-MS/MS conditions and separation parameters presented in Table 1–2.

**Table 1.** LC-MS/MS conditions for human blood plasma sample analysis.

|  | D- and L- |
| --- | --- |
| Scan Type: | MRM |
| Ion source: | Turbo spray |
| Probe position: | X=5.00, Y=5.2 |
| Polarity: | Positive |
| Run duration: | 11 min |
| Settling time (msec): | 0 |
| Pause time (msec): | 7.007 msec |
| Curtain gas: | 35 |
| CAD<sup>1</sup> gas: | 4 |
| ISV<sup>2</sup> (V): | 5000 |
| Temperature (°C): | 500 |
| Ion Source Gas 1 (GS 1): | 50 |
| Ion Source Gas 2 (GS 2): | 50 |
| Q1/Q3 resolution: | unit/unit |
| CEM<sup>3</sup> (V): | 2600 |

<sup>1</sup>Collision Gas (CAD)
<sup>2</sup>Ion Source Voltage (ISV)
<sup>3</sup>Channel Electron Multiplier (CEM)

**Table 2.** Inlet settings for human blood plasma D- and L- assay.

| Inlet Settings | D- and L- Assay |
| --- | --- |
| Analytical Column | Supelco Astek Chirobiotic V 250 x 2.1 mm, 5 µm |
| Guard Cartridge | None |
| Sample Temperature | 15 ± 5.0°C |
| Column Temperature | 30.0 ± 5.0°C |
| Mobile Phase A | Water:Acetic Acid:Ammonium Hydroxide: Methanol 5:1:0.3:993.5 |
| Mobile Phase B | N/A |
| Needle Rinse | Water:Acetic Acid:Ammonium Hydroxide: Methanol 5:1:0.3:993.5 |
| Flow Rate | 0.3 mL/min |
| Injection Volume | 10 µL |
| Run Time | 11 min |

## 3. Materials and Methods

### 3.1. Matrix Lot-to-Lot Comparison

Individual lots of human plasma differ according to a person’s overall health and collection efficiency [12]. A single lot of plasma is not enough to demonstrate the ruggedness of the assay system when such variability in the matrix exists [13]. Due to this, and in accordance with current CAP standards, a minimum of 10 lots of human matrix were collected from drug-free donors. These plasma samples were spiked at a low-level concentration with each analyte. These samples were prepared, extracted, and run as described above. The responses were calculated and the analyte to internal standard (IS) ratio and %CV is shown in Table 3.

**Table 3.** Matrix effects fortified with QC material to a concentration of 75 or 37.5 ng/mL and the %CV determined of the analyte/IS area ratio

| Drug / Metabolite | Mean | Matrix Comparison |
| --- | --- | --- |
|  | Analyte/IS Ratio | %CV Analyte/IS ratio |
| D-Amphetamine | 0.159 | 2.96 |
| L-Amphetamine | 0.17 | 3.26 |
| D-Methamphetamine | 0.376 | 4.22 |
| L-Methamphetamine | 0.433 | 4.48 |

### 3.2. Analytical Measurement Range

The analytical measurement range (AMR) of the assay refers to the concentration range that the assay is validated within and is determined by running a series of calibration curve standards covering a concentration range that encompass the concentration of analyte expected to find in patient samples [13]. The limits of the AMR were bounded by the lower limit of quantitation (LLOQ) and the upper limit of quantitation (ULOQ). The dynamic range may be described by a linear or quadratic fit [14,15]. Calibration curves were created using a minimum of six non-zero calibration points. To be accepted as the AMR, all points describing the calibration curve must pass within ± 20% of the nominal concentration [14]. Furthermore, the correlation coefficient (R^2^) for the calibration curve must be ≥ 0.99, or R should be ≥0.98 to be acceptable [16,17].

### 3.3. Sensitivity

The sensitivity of the assay system refers to the ability to reliably produce a signal throughout the entire calibration range, but specifically at the low-end of the calibration curve (the lower limit of quantitation, LLOQ) [18]. In hyphenated mass spectrometry assays, a signal that produces a signal to noise ratio (S/N) of ≥10 is considered valid for the LLOQ of an assay system [19]. Further, a S/N ratio of ≥5 is considered clear enough for the limit of detection. We test the sensitivity of the assay system by injecting six replicates of the LLOQ over three days and evaluating the resulting analytical determinations. Standard acceptance criteria of ±20% of nominal concentration apply.

### 3.4. Intra-day Precision and Accuracy

Intra-day precision and accuracy were determined using six replicates of each of three QC sample determinations and LLOQ from across at least three validation runs. Concentrations of the QC samples ranged across the curve, with the low QC set at approximately 3 times the LLOQ or less, the mid QC near the mid-range of the linear range of the curve, and the high QC set at 80-90% of the ULOQ. Percent accuracy and precision was determined for each individual measurement. To be accepted, the precision and accuracy for the replicate determinations must be ≤20% at each level.

### 3.5. Inter-day Precision and Accuracy

Inter-day precision and accuracy were determined using all replicates of each of three quality control (QC low, QC mid, and QC high) and LLOQ sample determinations from the analytical runs performed on 3 separate days. Concentrations of the QC samples ranged across the curve, with the low QC set around 3 times the LLOQ, the mid QC near the middle of the linear range, and the high QC set at 80-90% of the ULOQ. To be accepted, the precision and accuracy for the replicate determinations must be ≤20% at each level.

### 3.6. Exogenous Interfering Substances

Drugs that are known or suspected of interfering with similar bioanalytical systems should be evaluated to ensure that they do not suppress ionization or cause false-positive results for a given analyte [20,21]. The following medications were evaluated: over-the-counter mix, acetaminophen, ibuprofen, pseudoephedrine, caffeine, and naproxen. The following individual analytes were also tested: salicylic acid, phenylephrine, phentermine, diphenhydramine, and dextromethorphan. A high concentration of the possible interfering drug (typically 2,000 ng/mL or greater) was spiked into a low QC sample (15 – 75 ng/mL low QC). Acceptance criteria for a substance to be deemed as non-interfering is that the quantitated value for the low QC should be within ± 20% of the nominal value [21]. Furthermore, the spiked substance should not cause a false-positive or a false-negative result.

### 3.7. Partial Volumes and Dilutions

A spiked solution was created at a concentration above the ULOQ in this case 4000 ng/mL. The sample was run at discrete dilutions 1:5, 1:10, 1:20, and 1:50. Concentration determinations for all dilutions should be within ± 20% of the nominal value following correction for the dilution factor [22,23]. More recent literature suggests that the signal to noise ratio of both the quantification trace and the qualifying ion trace be 3-10 [24]. On occasion, an analyte will not have a quantifying ion that passes this criterion while still permitting the quantification trace to remain in a meaningful range. These instances should be documented in the laboratory SOP or validation report.

### 3.8. Carryover

Carryover is the presence of an analyte in a blank injection following a positive injection, resulting in a false-positive sample [25]. The injection needle should be washed in-between samples with a needle wash solution that is intended to remove contamination from the surface of the needle. The efficiency of this process is monitored during validation by assessing carryover in the following manner. Samples are injected in the following sequence: high QC, wash, high QC, wash, high QC, wash. Peak areas are integrated for both the analyte and internal standard. Peak area in the wash solutions should be 0.1 % or less of that found in the high QC standard. In addition, the mean of the peak area in the three wash solutions following the high QC replicates should be less than 20% of the LLOQ being used for the assay [26].

### 3.9. Additives and Clinical Conditions

Certain anticoagulants and slightly different matrices (plasma versus serum) can affect the performance of some assays. This is also true for conditions causing hemolytic, lipemic and icteric (high bilirubin) samples. Accordingly, we investigated these potential issues by comparing plasma with different anticoagulants including serum, plasma containing hemolyzed RBC, lipemic, and icteric plasma.

## 4. Results

### 4.1. Inter-day Average Back Calculated Calibration Standards

Table 4 shows the range of standard curves of the combined Amphetamine and the individual D- and Lanalytes and the correlation information. D- and Lcurve concentrations were half the above concentration ranging from 0.25 (neg) to 1000 ng/mL. Mean R values were all at least 0.99 indicating good fit to the data.

**Table 4.** Statistical analysis for each analyte standard curve over three assays.

| Drug / Metabolite | Curve | Mean R | RSD | Mean Slope | SD Slope | N | Fit |
| --- | --- | --- | --- | --- | --- | --- | --- |
|  | Range<br>(ng/mL) |  |  |  |  |  |  |
| D-Amphetamine | 2.5-1000 | 0.9998 | 0.00006 | 0.0042 | 0.0003 | 3.0000 | Quadratic |
| L-Amphetamine | 2.5-1000 | 0.9999 | 0.00012 | 0.0047 | 0.0003 | 3.0000 | Quadratic |
| D-Methamphetamine | 2.5-1000 | 0.9998 | 0.00006 | 0.0105 | 0.0011 | 3.0000 | Quadratic |
| L-Methamphetamine | 2.5-1000 | 0.9995 | 0.00010 | 0.0117 | 0.0018 | 3.0000 | Quadratic |

### 4.2. Accuracy and Precision, LLOQ

Six replicates of each validation level were run on at least three days. The D- and L-assay individually had an LLOQ of 2.5 ng/mL with a QC low of 7.5 ng/mL, a QC mid of 300 ng/mL and a high QC of 900 ng/mL. Tables 5–6 indicates mean, inter-assay and intra-assay statistic variability were all below 20%.

**Table 5.**
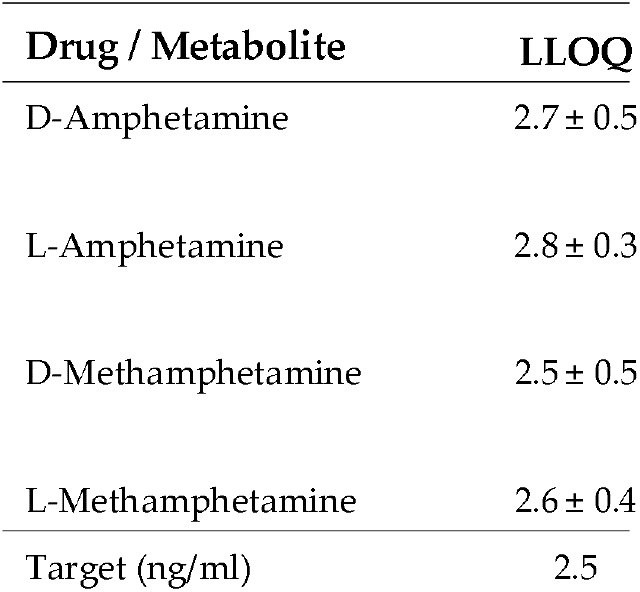
Inter-assay means and standard deviation (SD) of validation samples

**Table 6.** Inter-assay precision and accuracy over three (3) days with replicates of six (6) for each day

| Drug / Metabolite | %C |
| --- | --- |
|  | MIN |
| D-Amphetamine | 11.76 |
| L-Amphetamine | 8.02 |
| D-Methamphetamine | 12.53 |
| L-Methamphetamine | 7.14 |

### 4.3. Partial Volumes Accuracy and Precision

An MPA surrogate sample was prepared at 4000 ng/mL. To determine the concentration of this sample, a dilution must be made so the final concentration would be less than 2000 ng/mL to get it in the measurement range of the assay. Three replicates of four dilutions were made and tested: 1) 1:5 target 400 ng/mL; 2) 1:10 with a target of 200 ng/mL; 3) 1:20 with a target of 100 ng/mL; and 4) 1:50 with a target of 40 ng/mL. The results shown in Table 7 indicate that all analytes can be diluted at all levels.

**Table 7.** Dilution study: percent difference from expected with a 4000 (2000) ng/mL standard diluted as indicated.

| **Drug / Metabolite** | **1:5 Dilution** | **1:10 Dilution** | **1:20 Dilution** | **1: 100 Dilution** |
| --- | --- | --- | --- | --- |
| D-Amphetamine | -2.90 | 1.44 | -8.24 | -11.20 |
| L-Amphetamine | -4.40 | 1.28 | -7.28 | -9.96 |
| D-Methamphetamine | -2.84 | 0.32 | -9.04 | -10.08 |
| L-Methamphetamine | 1.44 | 1.32 | -8.16 | -8.76 |

### 4.4. Room Temperature, Refrigerator, and Freezer Stability

Samples with concentrations of 75, 800, or 1800 ng/mL were prepared in triplicate. One set was kept at room temperature (RT) overnight, a second set was kept in the refrigerator (RF) overnight and a third set was kept in the freezer (FZ) overnight. These validation samples were then run and compared to a triplicate preparation of QC samples that had been analyzed as normal. Table 8 show results less than 20% deviation from expected.

**Table 8.** stability testing QC samples were tested for stability after 3 freeze thaw cycles. They were also tested overnight at the indicated temperatures. A3 and 7 day post extraction study were also performed at 2-8 °C

| Drug / Metabolite | F/T 3 |
| --- | --- |
|  | QC |
| D-Amphetamine | QC 37.5 |
|  | QC 400 |
| L-Amphetamine | QC 37.5 |
|  | QC 400 |
| D-Methamphetamine | QC 37.5 |
|  | QC 400 |
| L-Methamphetamine | QC 37.5 |
|  | QC 400 |

### 4.5. Freeze-Thaw (FT) Stability

Validation samples with concentrations of 75, 800 or 1800 ng/mL were frozen at −20 °C and thawed in sequence with samples taken after each FT cycle for a maximum of 3 cycles. These validation samples were analyzed in triplicate and compared to a triplicate preparation of validation samples that had not been subjected to this FT cycle. The experimental results showed all meeting acceptance criteria.

### 4.6. Stability in Matrix

A series of triplicate samples were analyzed over 7 days for stability at room temperature, 4 °C and −20 °C. The results indicated that all analytes were stable for at least 7 days refrigerated and frozen. The analytes were stable at room temperature for 24h

### 4.7. Post Preperation Stability

A stability experiment was performed where samples were stored in the instrument (3 day) or refrigerator (7 day) and re-injected after 3 and 7 days. All samples were within 20% of the initial results.

### 4.8. Matrix Recovery and Matrix Effects

Table 3 indicates the effect of 10 different matrix lots tested by using a series of 37.5 ng/mL samples prepared in water, MPA and 10 different matrices. The results were acceptable with less than 20% CV across oral fluid, water and MPA meeting acceptance criteria. This is likely due to dilution in 1.5 mL Quantisal extraction buffer before extraction.

### 4.9. Selectivity

Multiple drugs that might have a potential for interfering with the assay analytes were run in the assay. Samples of 500 µL of 37.5 ng/mL QC were placed in a series of tubes to be run in triplicate. To the first set 50 µL of MeOH was added to act as the control. To the remaining tubes 50 µL of sample containing dextromethorphan, diphenhydramine, phenylephrine, salicylic acid, or combo (includes acetaminophen, caffeine, chlorpheniramine, ibuprofen, naproxen, and pseudoephedrine). These solutions were obtained from Cerilliant Corporation and were at a concentration of 1 mg/mL each except for the over-the-counter mix which was 100 µg/mL. Each solution was diluted to 20 µg/mL in methanol and this solution was used to spike samples as indicated above. Table 9 shows the results from this study. All samples met the acceptance criteria.

**Table 9.** Concomitant medications: The indicated medications prepared in methanol were spiked into a QC 37.5 standard and measured. The data indicates percent difference from a QC standard spiked with blank methanol at the same volume as the drug standards.

| Percent Difference from MEOH Spike |  |  |  |  |  |  |
| --- | --- | --- | --- | --- | --- | --- |
| Drug / Metabolite | Dextromethorphan | Phenylephrine | Diphenhydramine | Salicylic<br>Acid | Phentermine | OTC<br>Mix |
| D-Amphetamine | 9.31 | 0.76 | 1.79 | -3.35 | -1.83 | -0.76 |
| L-Amphetamine | 6.38 | 1.51 | 2.18 | -2.67 | -0.18 | -0.84 |
| D-Methamphetamine | 4.69 | -1.29 | -0.62 | -7.20 | -4.21 | -1.55 |
| L-Methamphetamine | 10.41 | -1.27 | 5.14 | -2.32 | -0.71 | 3.70 |

### 4.10. Additives and Clinical Conditions

Samples in triplicate at 37.5 ng/m were compared. These included serum defibrinated plasma, EDTA, and heparin. Clinical considerations included hemolytic, lipemic, and icteric samples. No deviations more than 15% was observed and only serum was more than 10% (Table 10). This indicates that none of these conditions adversely affects the measurement of these analytes.

**Table 10.** Alternate matrices, anti-coagulants, and disease states

| % Diff from Target |  |  |  |  |  |  |  |  |
| --- | --- | --- | --- | --- | --- | --- | --- | --- |
| Drug / Metabolite | Defibrinated | EDTA | Heparin | Serum | Hemolyzed | Lipemic | Icteric 2 | Icteric 20 |
| D-Amphetamine | -4.07 | -3.78 | -5.54 | -4.89 | -4.09 | -4.82 | -3.14 | -6.39 |
| L-Amphetamine | -2.88 | -2.55 | -5.04 | -5.58 | -5.08 | -4.94 | -3.54 | -6.13 |
| D-Methamphetamine | -4.52 | -7.00 | -9.06 | -12.28 | -10.18 | -7.43 | -7.04 | -8.64 |
| L-Methamphetamine | -4.60 | -5.40 | -6.61 | -11.16 | -10.70 | -8.05 | -7.37 | -7.15 |

## 5. Discussion

Urine and oral fluid are commonly preferred matrices for drug toxicological testing. On occasion, medical providers deem it medically necessary to order toxicology testing on a blood sample. Accordingly, this study describes a validated laboratory develop LC-MS/MS assay to quantify the enantiomeric forms of (meth)amphetamine) in human blood plasma samples. The utility of this assay is its concomitant use when (meth)amphetamine is detected in a broader confirmation panel in order to determine if the positive result was from the legal or illicit form of (meth)amphetamine. This assay removes the plausible deniability of illicit (meth)amphetamine use as an artifact of decongestant use or a false-positive due to prescriptive forms of amphetamine.

All analytes were well-behaved during development, providing a meaningful analytical measurement range, acceptable intra-day and inter-day precision and accuracy, specificity for the target analytes, and the assay demonstrated acceptable stability that allows for a reasonable laboratory workflow. The most important aspect of this assay was its specificity. It can reliably and definitively differentiate between the isomeric forms of (meth)amphetamine, as well as common decongestants and weight loss medication. Phentermine is a positional isomer of methamphetamine that laboratories need to ensure does not interfere in methamphetamine confirmation [27]. As a positional isomer, it shares a molecular weight and fragment pattern nearly indiscernible from methamphetamine by many LC-MS/MS methods. Chromatographic separation saw phentermine elute at 4.21 minutes, while methamphetamine eluted at 6.06 for the Disomer and 6.58 minutes for the L-isomer. Furthermore, phentermine only shared the qualifying ion signal with methamphetamine, but not the quantifying ion. Since the peaks were temporally separated by 1.85 minutes, phentermine shows no interference in the MRM window that is used for Dor Lmethamphetamine.

Physicians may choose blood as a matrix for toxicologic testing for numerous reasons [28]. While urine is the most frequently used biologic matrix, drug concentrations in urine do not necessarily reflect circulating concentrations of drugs. Oral fluids offer a secondary biological matrix, but there is poor penetration of certain drugs and their metabolites into the oral fluid compartment therefore limiting the utility of the assay [29]. Further, some individuals have difficulty providing a urine or oral fluid specimen and blood is consequently selected as a suitable sample for toxicology testing. The attractiveness of blood is that the drug levels detected are biologically available and both active and inactive metabolites are detectable, unlike in oral fluid. Likewise, unlike urine and oral fluid, it is exquisitely difficult to tamper with or adulterate blood samples [28].

This assay is reliable, reproducible, and removes plausible deniability from drug confirmations for D- and L(meth)amphetamine in blood plasma samples. It provides a fully quantitative analysis of each of the stereoisomers of two clinically significant stimulants—methamphetamine and its metabolite amphetamine, and it cleanly separates these analytical targets from interfering substances. The sensitivity of the assay is tremendous considering that this assay uses an older instrument (API SCIEX 4000) and can be meaningful for many clinical laboratories looking to perform drug confirmation studies for (meth)amphetamine.

## 6. Conclusion

The determination of prescription medications and illicit substances in human blood plasma is critical, notably where potential adulteration is a concern. Moreover, human blood plasma methods to quantitatively detect the enantiomeric forms of (meth)amphetamine has medical compliance and treatment implications [30], and warrants de-centralized offerings by independent clinical laboratories, especially in rural regions where (meth)amphetamine (mis)use is on the rise [31]. This paper describes a method to determine the D- and L- isomers of (meth)amphetamine using an older, less sensitive instrument (API SCIEX 4000) by using a liquid-liquid extraction method, concentration of the samples with a nitrogen dry-down, and a resuspension step. The method was validated in accordance with the US Food and Drug Administration and College of American Pathologists with an LLOQ of 2.5 ng/mL and ULOQ of 1000 ng/mL.

The novelty of this work is a sensitive method for the determination of the D- and L- isomers of (meth)amphetamine from an extract used for determination of 63 analytes from human K2EDTA plasma. The assay has good precision and accuracy and would a suitable addition to any clinical laboratory seeking confirmation of the enantiomeric forms of (meth)amphetamine following immunoassay or GC/MS positive results for methamphetamine or its metabolite amphetamine.

## Data Availability Statement

Harvard Dataverse https://doi.org/10.7910/DVN/TBEWWE

## References

1. Nichols, J. The epidemic within the pandemic: Behavioral health and substance use in the face of COVID-19. Pharm Today 2022, 28: 54–62. DOI: 10.1016/j.ptdy.2022.03.023

2. Naeim, M.; Rezaeisharif, A. Increased consumption of crystal (Methamphetamine) during the COVID-19 outbreak. Addictive Disorders & Their Treatment 2021, 20: 593–594. DOI: 10.1097/ADT.0000000000000250

3. Kiyokawa, M.; Cape, M.; Streltzer, J. Insights in public health: Methamphetamine use during COVID-19 in Hawaii. Hawai’i Journal of Health & Social Welfare 2021, 80: 117–118. DOI: 10.1056/NEJMra1511480. PMID: 26816013

4. Volkow, N. D. Collision of the COVID-19 and addiction epidemics. Annals of Internal Medicine 2020, 173: 61–62. DOI: 10.7326/M20-1212

5. Kuczenski, R.; Segal, D. S.; Cho, A. K.; Melega, W. Hippocampus norepinephrine, caudate dopamine and serotonin, and behavioral responses to the stereoisomers of amphetamine and methamphetamine. Journal of Neuroscience 1995, 15: 1308–1317. DOI: 10.1523/JNEUROSCI.15-02-01308.1995

6. Mendelson, J.; Uemura, N.; Harris, D.; Nath, R. P.; Fernandez, E.; Jacob III, P.; Jones, R. T. Human pharmacology of the methamphetamine stereoisomers. Clinical Pharmacology & Therapeutics 2006, 80: 403–420. DOI: 10.1016/j.clpt.2006.06.013

7. Bordoloi, M.; Chandrashekar, G.; Yarasi, N. ADHD in adults and its relation with methamphetamine use: national data. Current Developmental Disorders Report 2019, 6: 224–227. DOI: 10.1007/s40474-019-00174-w

8. Peters, F. T.; Meyer, M. R. Analytical techniques for the detection of novel psychoactive substances and their metabolites. In Novel Psychoactive Substances, Peters, F. T., Meyer, M. R., Eds.; Academic Press: San Diego, California, USA, 2013; pp. 131–157.

9. Robbins, B., Carpenter, R. E., Long, M. & Perry, J. A human oral fluid assay for D- and L-isomer detection of amphetamine and methamphetamine using liquid-liquid extraction. Journal of Analytical Methods in Chemistry (in press).

10. Peters, F. T.; Schaefer, S.; Staack, R. F.; Kraemer, T.; Maurer, H. H. Screening for and validated quantification of amphetamines and of amphetamine-and piperazine-derived designer drugs in human blood plasma by gas chromatography/mass spectrometry. Journal of Mass Spectrometry 2003, 38: 659–676. DOI 10.1002/jms.483

11. Robbins, B.; Carpenter, R. E.; Long, M.; Perry, J. Quantifying 64 drugs, illicit substances, and D- and L- isomers in human oral fluid with liquid-liquid extraction. BioRxiv 2022, DOI: 10.1101/2022.10.29.514362

12. Caporossi, L.; Santoro, A.; Papaleo, B. Saliva as an analytical matrix: State of the art and application for biomonitoring. Biomarkers 2010, 15: 475–487. DOI: 10.3109/1354750X.2010.481364

13. Gjerde, H.; Langel, K.; Favretto, D.; Verstraete, A.G. Detection of illicit drugs in oral fluid from drivers as biomarker for drugs in blood. Forensic Sci International 2015, 256: 42–45. DOI: 10.1016/j.forsciint.2015.06.027

14. Killeen, A.A.; Long, T.; Souers, R.; Styer, P.; Ventura, C.B.; Klee, G.G. Verifying performance characteristics of quantitative analytical systems: calibration verification, linearity, and analytical measurement range. Arch of Path & Lab Med. 2014, 138: 1173–1181. DOI: 10.5858/arpa.2013-0051-CP

15. Rule, G. S.; Clark, Z. D.; Yue, B.; Rockwood, A.L. Correction for isotopic interferences between analyte and internal standard in quantitative mass spectrometry by a nonlinear calibration function. Anal Chem 2013, 85: 3879–3885. DOI: 10.1021/ac303096w

16. Asuero, A.G.; Sayago, A.; González, A.G. The correlation coefficient: an overview. Critical Rev in Anal Chem 2006, 36: 41–59. DOI: 10.1080/10408340500526766

17. Van Loco, J.; Elskens, M.; Croux, C.; Beernaert, H. Linearity of calibration curves: use and misuse of the correlation coefficient. Accreditation and Qual Assurance 2002, 7: 281–285. DOI: 10.1007/s00769-002-0487-6

18. Liang, H. R.; Foltz, R. L.; Meng, M.; Bennett, P. Ionization enhancement in atmospheric pressure chemical ionization and suppression in electrospray ionization between target drugs and stable-isotope-labeled internal standards in quantitative liquid chromatography/tandem mass spectrometry. Rapid Comm in Mass Spect 2003, 17: 2815-2821. DOI 10.1002/rcm.1268

19. Meyer, C.; Seiler, P.; Bies, C.; Cianciulli, C.; Wätzig, H.; Meyer, V. R. Minimum required signal-to-noise ratio for optimal precision in HPLC and CE. Electrophoresis 2012, 33: 1509–1516. DOI 10.1002/elps.201100694

20. Gosetti, F.; Mazzucco, E.; Zampieri, D.; Gennaro, M.C. Signal suppression/enhancement in high-performance liquid chromatography tandem mass spectrometry. J of Chromatography A 2010, 1217: 3929–3937. DOI: 10.1016/j.chroma.2009.11.060

21. Peters, F.T.; Remane, D. Aspects of matrix effects in applications of liquid chromatography–mass spectrometry to forensic and clinical toxicology—a review. Anal and Bioanal Chem 2012, 403: 2155–2172. DOI: 10.1007/s00216-012-6035-2

22. Viswanathan, C. T.; Bansal, S.; Booth, B.; DeStefano, A. J.; Rose, M. J.; Sailstad, J.; Weiner, R. Quantitative bioanalytical methods validation and implementation: best practices for chromatographic and ligand binding assays. Pharma Research 2007, 24: 1962–1973. DOI: 10.1007/s11095-007-9291-7

23. Mannu, R. S.; Turpin, P.E.; Goodwin, L. Alternative strategies for mass spectrometer-based sample dilution of bioanalytical samples, with particular reference to DBS and plasma analysis. Bioanalysis 2014, 6: 773–784. DOI: 10.4155/bio.13.320

24. Cai, H. L.; Wang, F.; Peng, W. X.; Zhu, R. H.; Deng, Y.; Jiang, P.; Chen, C. Quantitative analysis of erythromycylamine in human plasma by liquid chromatography-tandem mass spectrometry and its application in a bioequivalence study of dirithromycin enteric-coated tablets with a special focus on the fragmentation pattern and carryover effect. J of Chromatography B 2014, 947:156–163. DOI: 10.1016/j.jchromb.2013.12.019

25. Food and Drug Administration. Guidance for Industry: Bioanalytical Method Validation, US Department of Health and Human Services, FDA, Center for Drug Evaluation and Research, Rockville, MD, 2001.

26. Food and Drug Administration. Guidance for Industry: Bioanalytical Method Validation, US Department of Health and Human Services, FDA, Center for Drug Evaluation and Research, Rockville, MD, 2018.

27. Kranenburg, R. F.; Lukken, C. K.; Schoenmakers, P. J.; van Asten, A. C. Spotting isomer mixtures in forensic illicit drug casework with GC–VUV using automated coelution detection and spectral deconvolution. Journal of Chromatography B 2021: 1173: 1–10. DOI: 10.1016/j.jchromb.2021.122675

28. de Campos, E. G.; da Costa, B. R. B.; dos Santos, F. S.; Monedeiro, F.; Alves, M. N. R.; Santos Junior, W. J. R.; De Martinis, B. S. Alternative matrices in forensic toxicology: A critical review. Forensic Toxicology 2021, 1–18. DOI: 10.1007/s11419-021-00596-5

29. Patrick, M.; Parmiter, S.; Mahmoud, S. H. Feasibility of using oral fluid for therapeutic drug monitoring of antiepileptic drugs. European Journal of Drug Metabolism and Pharmacokinetics 2021, 46: 205–223. DOI: 10.1007/s13318-020-00661-1

30. Carpenter, R. E.; Silberman, D. Takemoto, J.K. Transforming prescription opioid practices in primary care with change theory. Health Services Insights 2021, 14: 1–8. DOI: 10.1177/11786329211058283

31. Hansen, E. R.; Carvalho, S.; McDonald, M.; Havens, J. R. A qualitative examination of recent increases in methamphetamine use in a cohort of rural people who use drugs. Drug and Alcohol Dependence 2021, 229. DOI: 10.1016/j.drugalcdep.2021.109145

